# Terminally Differentiated Influenza-Specific Effector Memory B Cells Circulate after Live Attenuated Influenza Vaccination

**DOI:** 10.64898/2026.01.17.700068

**Authors:** Anoma Nellore, Christopher Fucile, Christopher Scharer, Jack T Geer, Jacey Lemonds, Betty Mousseau, Fen Zhou, Esther Zumaquero, Jobaida Akhter, Ellie Ivanova, Page Murray, Valeria Estrada Navarro, Paul A Goepfert, Ralf Duerr, Ramin S Herati, Alexander Rosenberg, Frances E Lund, Troy Randall

## Abstract

Live attenuated influenza vaccination (LAIV) is the only FDA approved mucosal vaccine. Easily assayed, circulating correlates of protection after LAIV are lacking. Using fluorochrome labeled hemagglutinin (HA) antigen, we previously identified a subset of HA-specific (HA^+^) IgD^neg^ memory B cells that circulate after inactivated intramuscular vaccination (IIV), express the master transcriptional regulator, T-bet, as well as effector memory genes and predict durable antibody (Ab) responses to IIV. Here we profile the circulating HA^+^ IgD^neg^ memory B cell response in a cohort of immunized patients who seroconvert after LAIV to identify which, if any, circulating HA^+^ B cells predict antibody responses after LAIV. Although we report LAIV elicits circulating T-bet^+^ HA^+^ IgD^neg^ B cells that are phenotypically similar to those we described after IIV, we find no correlation between the magnitude of these cells and systemic HA-IgG responses after LAIV. Supervised and unsupervised analyses demonstrate that unlike IIV, LAIV preferentially elicits circulating HA^+^ IgD^neg^ B cells that co-express *TBX21* and the terminal effector cell gene, *Zeb2*. Consistent with their terminal differentiation status, LAIV-elicited T-bet^+^ cells cannot be recalled as short-lived antibody-secreting cells (ASCs) after systemic or mucosal antigen re-challenge. We conclude that the transcriptional profiles and functions of HA^+^ IgD^neg^ B cells vary by influenza vaccine platform.

## Introduction

Influenza is a significant global threat and causes 290,000 to 650,000 annual fatalities world-wide.^1^ As an RNA virus, influenza continuously evolves via antigenic shift or drift to result in seasonal or pandemic spread of disease. Influenza vaccines are formulated as intramuscular and inactivated (IIV) or as live attenuated (LAIV) for intranasal mucosal delivery. Vaccines act by inducing antigen-specific humoral immune responses to immunogenic influenza proteins including hemagglutinin (HA). In the context of influenza vaccination of adults, who are almost certainly flu experienced, this antigen-specific vaccine response is driven by expansion of pre-existing memory B cell populations. Some of these memory B cells are recalled into germinal centers to interact with memory T follicular helper cells (Tfh) and to produce daughter memory and antibody-secreting cells (ASCs), which can mediate long-term protection. Memory B cells can be further classified into (1) cells that have stem-like properties and can be recalled into germinal center responses to generate daughter memory B cells and ASCs and (2) cells with effector properties as poised to quickly and terminally differentiate into plasmablasts (PBs) or (3) effector cells that directly secrete cytokines.^2, 3^

Mucosally delivered influenza viral antigen has been shown in murine models and among immunized humans to elicit mucosal, tissue resident memory B cells as critical mediators of first-line defense against challenge infections.^4, 5, 6^ However, the efficacy of LAIV versus IIV in humans is inconsistent. Historically, in pediatric cohorts, LAIV has been shown to elicit superior immunogenicity than IIV.^7^ In contrast, in adults, LAIV is not uniformly protective.^8^ In fact, during the 2015-2016 influenza season, LAIV was removed from the US market because of a lack of efficacy, even among pediatric patients, against the H1N1 strain.^9^ Importantly, no easily assayed circulating correlate for immune protection after LAIV has been defined and this limits our ability to understand the quality of immune protection after LAIV.^10^

Until recently, the only antigen-specific outputs that could be assayed after IIV were PBs and circulating Tfh (cTfh) as measures of IIV-responsiveness.^11, 12, 13^ Now fluorochrome labeled influenza HA antigen probes or tetramers have been utilized as reagents that permit the direct study of antigen-specific B cells in humans that can circulate at various time points after vaccination. Using these tetramer reagents, the circulating IIV-elicited HA-specific (HA^+^) B cell compartment in adults has been defined as mutated, consistent with the properties of memory B cells.^14, 15, 16^ These HA^+^ memory B cells have also been described as functionally heterogeneous. For example, we and others have classified IIV-elicited circulating HA^+^ memory B cells according to expression of the lineage defining effector transcription factor, T-bet, as well as atypical memory markers, CD21, FcRL5 and CD11c. T-bet^+^ HA^+^ memory B cells circulate as early as 7 days after IIV, are identified in the blood as late as 60-120 days after vaccination, and predict durable HA IgG immune responses to IIV. T-bet^+^ HA^+^ memory B cells are transcriptionally programmed as effector memory and comprise clones that are preferentially recalled as PBs after antigen re-challenge for immediate immune protection from disease. Thus, circulating T-bet^+^ HA^+^ memory B cells represent a candidate rapid, easily assayed biomarker of vaccine response but whether this readout generalizes across other vaccine platforms, like LAIV, is not completely known. ^14, 15, 16^

Here we apply fluorochrome labeled B cell tetramers to evaluate HA-specific (HA^+^) memory B cells in the blood after LAIV in cohort of inoculated healthy adults who sero-covert after LAIV. We specifically compare the functional properties of HA-specific (HA^+^) memory B cells that circulate after LAIV to those that circulate after IIV in order to assess which, if any, circulating memory B cells can function as candidate correlates of durable immune protection. We find that we can identify T-bet^+^ HA^+^ memory B cells that circulate after LAIV and are phenotypically similar to those in circulation after IIV. However, unlike our findings in subjects after IIV, we find that T-bet^+^ HA^+^ memory B cells do not correlate with durable Ab after LAIV. Bulk cell and single cell analyses demonstrate that LAIV-elicited T-bet^+^ HA^+^ memory B cells are transcriptionally unique from IIV-elicited counterparts by upregulating terminal effector cell genes. We additionally find that circulating LAIV-elicited T-bet^+^ HA^+^ memory B cells cannot be recalled as PBs after antigen re-challenge, consistent with their terminal effector cell differentiation state. Together, these data suggest heterogeneity in circulating T-bet^+^ B cell functions by influenza vaccine platform.

## Results

### HA^+^ IgD^neg^ B cells circulate in adults who seroconvert after LAIV

During the 2015-2016 influenza vaccine season, we enrolled 36 healthy subjects to receive either LAIV (N = 17) or IIV (N = 19) with serial blood draws after vaccination. Subjects self-reported as missing the prior season influenza vaccination and being immunologically healthy. The LAIV cohort enrolled a higher proportion of women than the IIV cohort but the two cohorts did not differ according to other demographics, including age and ethnicity [Supplemental Table 1].

To characterize the immune response to these vaccines, we enumerated the frequencies of known early (day 7) circulating cellular markers of vaccine response, namely CD27^+^CD38^+^ short-lived antibody-secreting cells (ASCs), also called plasmablasts (PBs)^11, 12^ and CXCR5^+^PD1^+^ CD4^+^ T follicular helper (cTfh) cells.^13, 17, 18^ Gating strategies are shown in Supplemental Figure 1A-B. We found an increase in the circulating PB [Fig 1A-B] and cTfh cell fractions [Fig 1D-E] after IIV and LAIV. However, we found that the frequency of circulating LAIV-elicited PBs and LAIV-elicited cTfh cells to be significantly lower (p <0.0001) than IIV-elicited PBs and IIV-elicited cTfh cells respectively [Fig 1C, F].

**Figure 1.**
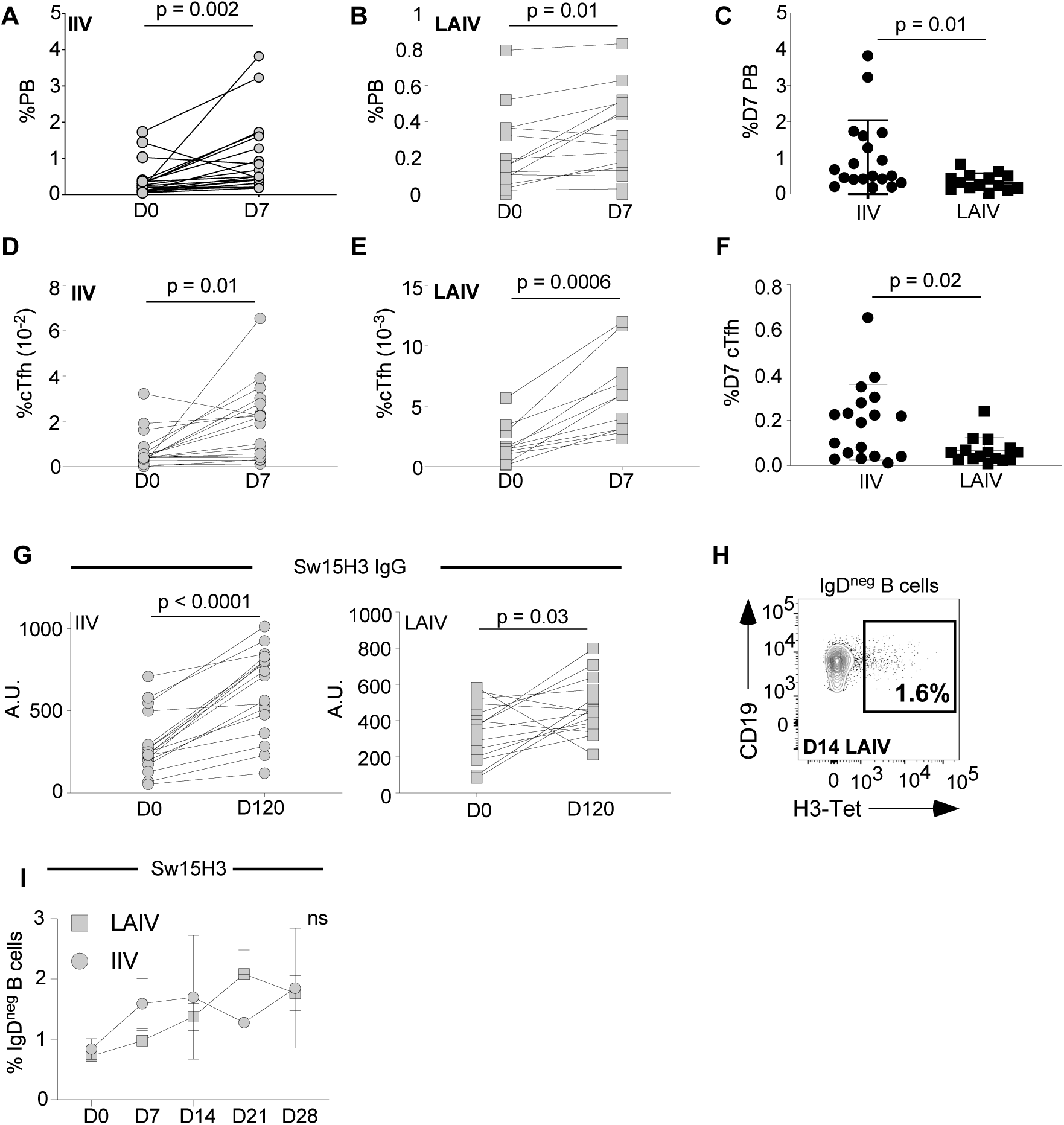
HA B cell tetramers identify circulating HA^+^ IgD^neg^ B cells after LAIV. Nineteen healthy subjects were immunized with the 2015-2016 IIV and seventeen healthy subjects were immunized with the 2015-2016 LAIV (see Supplemental Table 1 for demographics). Peripheral blood mononuclear cells (PBMCs) were drawn at weekly timepoints after inoculation. **(A)** Percentage of plasmablasts (PBs) from total live CD19^+^ B cells (see Supplemental Fig 1A for gating strategy) that circulate at day 0 and day 7 after IIV inoculation. **(B)** Percentage of PBs from total live CD19^+^ B cells that circulate at day 0 and day 7 after LAIV inoculation. **(C)** Dot plot depicting the percent PBs that circulate after LAIV (square) versus IIV (circle) **(D)** Percentage of circulating T follicular helper cells (cTfh) from total live CD4^+^ T cells (see Supplemental Fig 1B for gating strategy) at day 0 and day 7 after IIV inoculation **(E)** Percentage of cTfh cells from total live CD4^+^ T cells at day 0 and day 7 after LAIV inoculation **(F)** Dot plot depicting the percent cTfh cells that circulate after LAIV (square) versus IIV (circle) **(G)** IgG antibody titer (AU) against the Switzerland H3N2 strain in IIV and LAIV inoculated subjects **(H)** FACS plot gated on live IgD^neg^ B cells 14 days after LAIV depicts H3-tetramer binding (H3^+^) B cells (see Supplemental Fig 1C for gating strategy). **(I)** Percent of Sw15-H3-binding (H3^+^) IgD^neg^ B cells that circulate after IIV (circle) and LAIV (square) inoculation.

Fluorochrome labeled influenza (hemagglutinin or HA) antigen or HA B cell tetramers have been used to identify HA-specific (HA^+^) B cells in the blood after IIV.^14, 15, 16^ Therefore, we hypothesized that HA^+^ B cells might also circulate after LAIV. To test this, we used vaccine antigen-matched HA tetramers to enumerate HA^+^ IgD^neg^ B cells [Gating Strategy, Suppl Fig 1A] in the blood within one month of LAIV. The Ca09 H1N1 antigen was reported as poorly immunogenic in the 2015-2016 LAIV^9^ and in keeping with this finding, subjects immunized with 2015-2016 LAIV did not mount a durable day 120 Ca09 H1 IgG response after LAIV [Supplemental Fig 1 C, D] nor did they have an appreciable increase in the frequency of Ca09H1^+^ IgD^neg^ B cells in blood relative to participants who received IIV [Supplemental Fig 1E]. In contrast, we found our 2015-2016 LAIV inoculated cohort and 2015-2016 IIV inoculated cohort both mounted a durable day 120 H3 IgG response to the Sw15 H3N2 vaccine antigen [Fig 1G]. We also found appreciable numbers of H3^+^ IgD^neg^ B cells in the blood after 2015-2016 LAIV [Fig 1H]. Moreover, the frequencies of LAIV-elicited versus IIV-elicited H3^+^ IgD^neg^ B cells were similar at weekly timepoints within one month of inoculation [Fig 1I]. These data suggest that circulating HA^+^ IgD^neg^ B cells may be more reliably observed as circulating LAIV-elicited cellular outputs over other known vaccine antigen-specific cellular outputs, like cTfh and PBs.

### Supervised Analyses Demonstrate LAIV and IIV Elicit Transcriptionally Heterogeneous HA^+^ IgD^neg^ B Cells by T-bet expression status

We and others have reported that antigen-specific B cells that circulate after IIV are transcriptionally and functionally heterogenous by expression of the lineage defining transcription factor, T-bet.^14, 15, 16^ We additionally report that the magnitude of IIV-elicited T-bet^+^ HA^+^ IgD^neg^ B cells predicts durable HA IgG Ab 120 days after immunization. Similar to our observations from IIV-inoculated individuals, we found H3^+^ IgD^neg^ B cells that circulate in subjects after 2015-2016 LAIV could also be classed into T-bet^+^ and T-bet^neg^ subsets that expand over time in the blood [Supplemental Fig 2A-C]. We tested for correlations between day 120 H3 IgG Ab and known antigen-specific subsets (cTfh and PBs) and found no significant association [Supplemental Fig 2D-E]. Next, we tested for correlations between the magnitude of either the day 14 or day 21 T-bet^+^ or T-bet^neg^ HA^+^ IgD^neg^ subsets and day 120 H3 IgG Ab. We chose the day 14 time point because this is when the frequency of LAIV-elicited HA^+^ IgD^neg^ B cells starts to increase in blood and the day 21 time point because this is when the frequency of LAIV-elicited HA^+^ IgD^neg^ B cells is at its peak [Fig 1I]. We did not identify an association between the magnitude of either the day 14 [Fig 2A-B] or day 21 [Fig 2A-B] T-bet^+^ or T-bet^neg^ HA^+^ IgD^neg^ subsets and day 120 H3 IgG Ab. This is unlike our previously reported findings that T-bet^+^ HA^+^ IgD^neg^ B cells that circulate after IIV predict IIV-elicited day 120 HA IgG. Therefore, we speculated that LAIV-elicited and IIV-elicited T-bet^+^ B cells may be functionally dissimilar.

**Figure 2.**
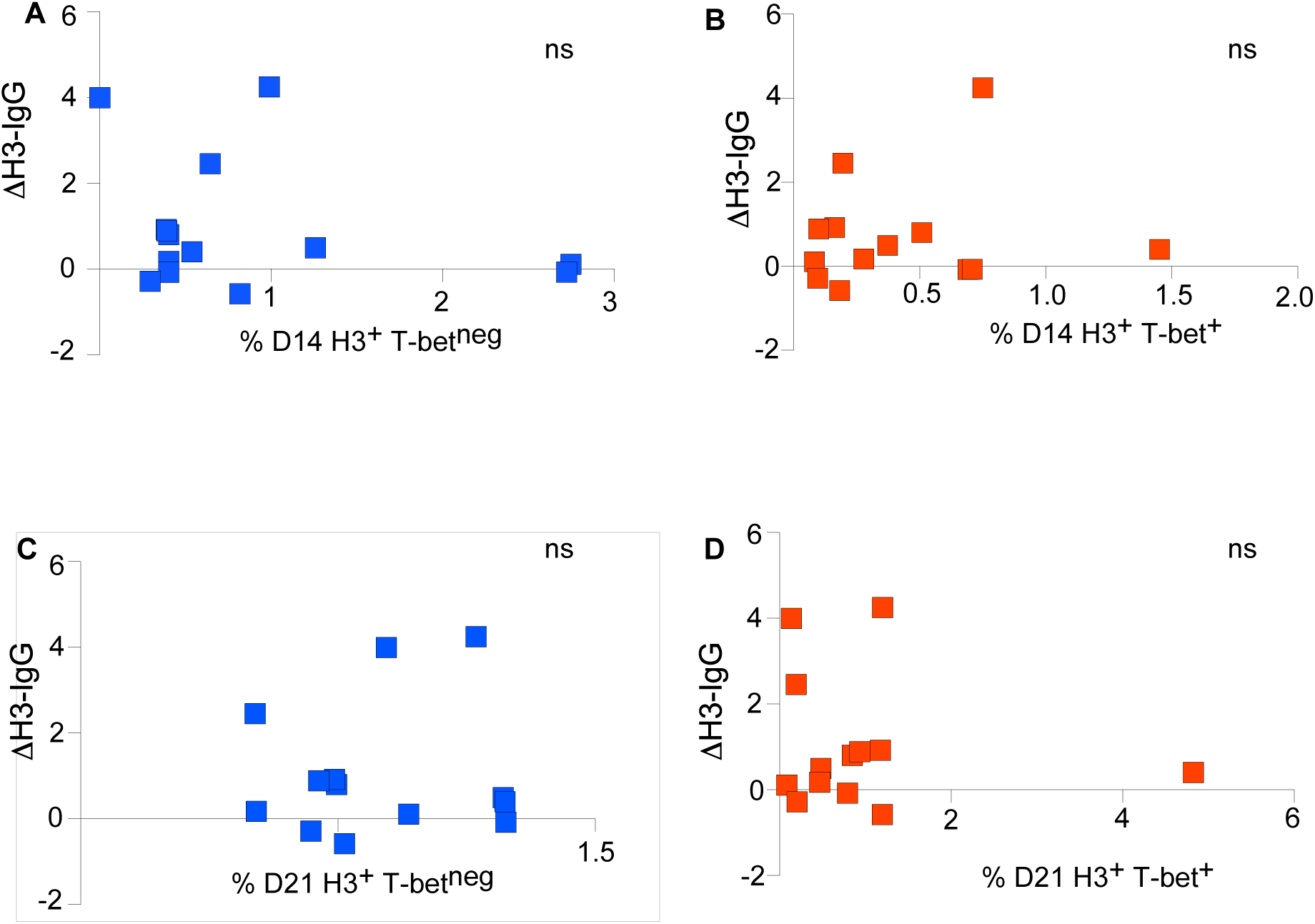
Circulating T-bet^+^ and T-bet^neg^ H3^+^ IgD^neg^ B cells do not predict durable H3 antibody response. As described in Figure 1, seventeen healthy subjects received 2015-2016 LAIV. Correlation between the fold change in H3-IgG Ab at days 0 and 120 and the magnitude of **(A, C)** H3^+^T-bet^neg^ IgD^neg^ B cells or **(B, D)** H3^+^T-bet^+^ IgD^neg^ B cells at day 14 (**A, B**) or day 21 (**C, D**) from the seventeen LAIV inoculated subjects. Gating strategies for H3^+^ T-bet^+^ IgD^neg^ B cells and H3^+^ T-bet^neg^ IgD^neg^ B cells are shown in Supplemental Fig 2A-C.

To start to test this hypothesis, we used an established B cell surface marker surrogate of T-bet expression, FcRL5,^16^ to sort purify T-bet^+^ HA (H3)^+^ IgD^neg^ B cells (T-bet^+^) and T-bet^neg^ HA (H3)^+^ IgD^neg^ B cells (T-bet^neg^) that circulated 14 days after 2015-2016 LAIV for bulk RNA-seq. We found 273 differentially expressed genes (DEGs). [Fig 3A] DEGs that are up-regulated in T-bet^+^ over T-bet^neg^ cells include effector genes, *TBX21*, *ZEB2*, *BATF* and DEGs that are down-regulated in T-bet^+^ over T-bet^neg^ cells include markers of lymph node homing, *CXCR5*.

**Figure 3.**
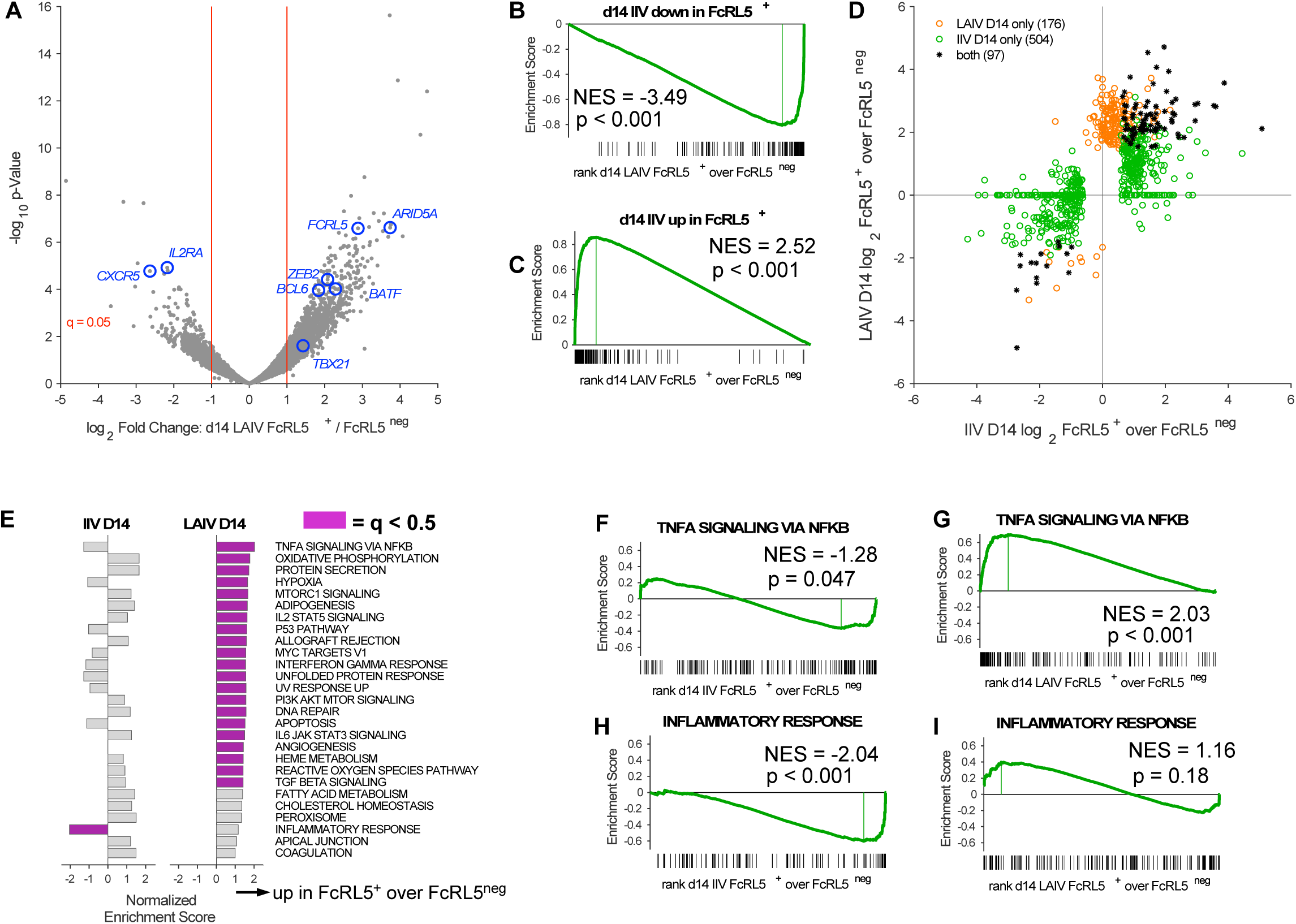
Bulk RNA-seq demonstrate unique transcriptional signature of LAIV versus IIV elicited HA^+^ IgD^neg^ B cells by FcRL5 expression status. FcRL5^+^ HA^+^ IgD^neg^ B cells and FcRL5^neg^ HA^+^ IgD^neg^ B cells were sort purified from N = 5 LAIV immunized subjects at day 14 or bulk RNA-seq. **(A)** Volcano plot depicts differentially expressed genes (DEGs). Genes of interest are annotated in blue. Threshold of q value at 0.05 is annotated by red line. **(B-C)** GSEA comparing the ranked gene list from LAIV-elicited H3^+^ FcRL5^+^ IgD^neg^ B cells over H3^+^ FcRL5^neg^ IgD^neg^ B cells that are either downregulated **(B)** or upregulated **(C)** in IIV-elicited H1-specific FcRL5^+^ IgD^neg^ B cells over H1-specific FcRL5^neg^ IgD^neg^ B cells (ref 16) **(D)** Scatterplot depicts the fold change of differentially expressed genes in LAIV versus IIV elicited FcRL5^+^ over FcRL5^neg^ subsets. Genes are colorized by assignment to IIV dataset (green), LAIV dataset (orange) or both datasets (black). **(E)** Bar plot of Normalized Enrichment Score for each Hallmark gene set is shown. Significant bar plots (q <0.05) are colorized purple. **(F-I)** GSEA comparing ranked gene lists from IIV-elicited FcRL5^+^ over FcRL5^neg^ subsets **(F, H)** LAIV elicited FcRL5^+^ over FcRL5^neg^ subsets **(G, I)** to Hallmark gene sets TNF-α signaling via NF-κb **(F-G)** and Inflammatory response **(H-I)**.

Next, we used Gene Set Enrichment Analyses (GSEA)^19^ to compare the transcriptome of FcRL5^+^ over FcRL5^neg^ subsets elicited 14 days after LAIV to the previously published transcriptome of FcRL5^+^ over FcRL5^neg^ subsets elicited 14 days after IIV.^16^ We found an enrichment in genes up-regulated or down-regulated by LAIV-elicited and IIV-elicited FcRL5^+^ cells in comparison to LAIV and IIV-elicited FcRL5^neg^ cells respectively. [Fig 3B-C]. However, when we evaluated the numbers of DEGs shared between FcRL5^+^ over FcRL5^neg^ subsets from LAIV inoculated versus IIV inoculated subjects, we found shared DEGs to be in the minority (97 genes) and the majority of DEGs (176 genes) were found as unique to either the LAIV or IIV comparisons [Fig 3D]. These data suggest that the functional properties of FcRL5^+^ and FcRL5^neg^ subsets may differ by vaccine platform.

To start to test for functional differences in FcRL5^+^ and FcRL5^neg^ subsets by vaccine platform, we performed GSEA of the FcRL5^+^ over FcRL5^neg^ comparison in LAIV inoculated subjects against curated MSigDB Hallmark gene sets. We found 27 Hallmark datasets as positively enriched with an FDR < 0.05. No Hallmark gene sets were found as negatively enriched [Fig 3E]. Next, we tested whether these 27 Hallmark gene sets were also enriched against IIV-elicited FcRL5^+^ over FcRL5^neg^ subsets [Fig 3E]. We found that day 14 LAIV-elicited and day 14 IIV-elicited FcRL5^+^ over FcRL5^neg^ subsets shared positive enrichment against curated Hallmark “Oxidative Phosphorylation”, “MTORC1 Signaling”, “Reactive oxygen species” and “IL-6-JAK-STAT3” signaling gene sets. However, we also found certain Hallmark data sets as unique to LAIV-elicited versus IIV-elicited subsets [Fig 3E]. For example, the Hallmark gene set “TNF-α signaling via NF-κb” was significantly and positively enriched against LAIV-elicited day 14 FcRL5^+^ over FcRL5^neg^ subsets but significantly and negatively enriched against day 14 IIV-elicited and day 7 FcRL5^+^ over FcRL5^neg^ subsets [Fig 3F-G]. The Hallmark dataset “Inflammatory Response” was significantly and positively enriched against LAIV-elicited day 14 FcRL5^+^ over FcRL5^neg^ subsets but significantly and negatively enriched against IIV-elicited day 14 FcRL5^+^ over FcRL5^neg^ HA^+^ B cells [Fig 3H-I]. Since IIV-elicited HA^+^IgD^neg^ B cells also circulate at scale within 7 days of immunization [Fig 1I], we wanted to ensure that our timepoint of comparison to IIV-elicited cells did not affect our comparison. Therefore, we evaluated published transcriptional data from day 7 IIV-elicited FcRL5^+^ over FcRL5^neg^ HA^+^ IgD^neg^ B cells.^16^ While the polarity or significance of certain Hallmark pathways were not conserved (e.g. hypoxia and oxidative phosphorylation) across day 7 and day 14 comparisons, we still identified a significant and negative enrichment against the Hallmark “Inflammatory Response” data set and a negative enrichment against the Hallmark “TNF-α signaling via NF-κb” in day 7 comparisons [Supplemental Fig 3A]. Collectively, these analyses suggest that relative to their T-bet^neg^ counterparts, T-bet^+^ HA^+^ IgD^neg^ B cells that circulate after LAIV exhibit different transcriptional properties from those that circulate after IIV.

### Unsupervised Analyses Demonstrate LAIV and IIV Elicit Transcriptionally and Clonotypically Heterogeneous HA^+^ IgD^neg^ B Cells with Effector and Stem Cell Programs

Our bulk RNA-seq data raise the possibility that LAIV and IIV elicit heterogeneous circulating HA^+^ IgD^neg^ B cell subsets. To test this possibility more rigorously and in an unsupervised manner, we administered the 2018-2019 LAIV (n =3) or the 2019-2020 LAIV (n =3) to N = 6 healthy subjects and the 2018-2019 IIV (n=2) or 2019-2020 IIV (n=2) to N = 4 healthy subjects. We sort-purified vaccine antigen-matched HA^+^ (H1 or H3) cells that circulated within one month after vaccination for single cell gene expression and matched BCR sequencing. We integrated the gene expression from all inoculated subjects with Seurat^20^, making note to exclude any BCR heavy and light chain gene segments and used Harmony^21^ to correct for batch effects. After filtering out cells with high mitochondrial DNA (including PB clusters 6 and 11, *not shown*), we resolved gene expression from 13902 total HA-binding IgD^neg^ B cells for single cell analyses. We resolved VDJ calls from 34% of HA-binding IgD^neg^ B cells for B cell receptor (BCR) repertoire analyses.

With this approach, we identified 9 transcriptionally discrete clusters based on gene expression patterns [Fig 4A]. Cell numbers per cluster are depicted in Supplemental Fig 4A. The 9 transcriptionally discrete clusters could be identified in 9 out of 10 donors tested; the donor lacking all 9 transcriptionally discrete clusters had the least cells amplified [Supplemental Fig 4B]. We used the ROGUE scoring method^22^ to evaluate how similar the transcriptomes within a cluster are to each other and we found all clusters to have a score of greater than 0.9, suggesting the relative purity of transcriptional expression in each of our resolved clusters [Supplemental Fig 4C].

**Figure 4.**
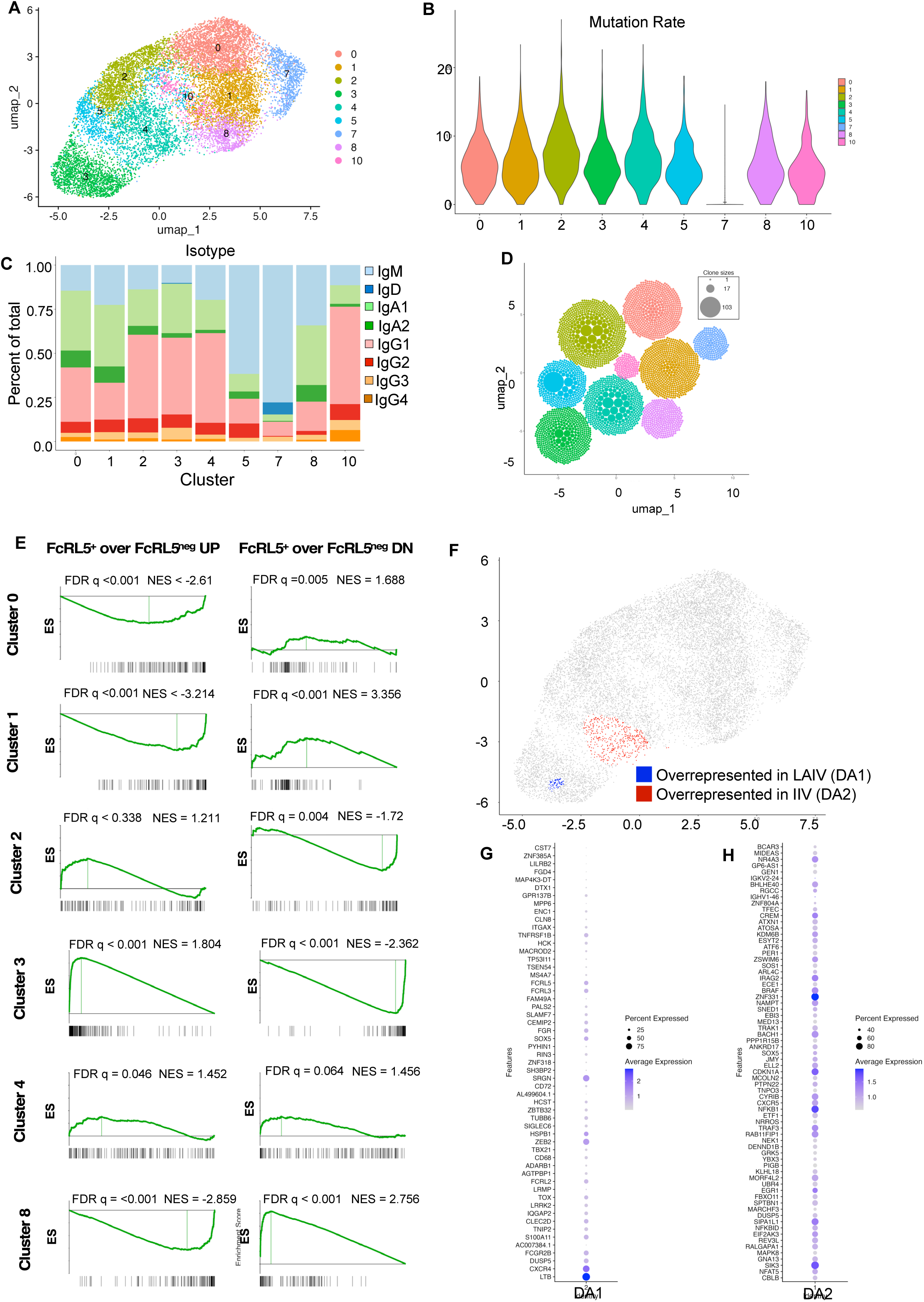
Single cell sequencing identifies differentially abundant HA^+^ IgD^neg^ B cells by vaccine platform. Six individuals received LAIV and 5 individuals received IIV. Vaccine antigen-matched HA^+^ IgD^neg^ B cells were sort-purified for single cell sequencing and data was integrated using Harmony (ref 21). **(A)** UMAP projection of HA^+^ IgD^neg^ B cells clustered by transcriptional similarity. (Supplemental Figure 4A shows total cell number by cluster. Supplemental Figure 4B shows cellular contributions to clusters by individual.) **(B)** Violin plot depicts IGHV mutation rates for clusters. **(C)** Bar plot depicts isotype frequency by cluster assignment. **(D)** Cells were annotated using the Monaco Immune Database. Bar plot depicts frequency of annotated cells by cluster. **(E)** UMAP projection of clonality by cluster (see Figure 4A for cluster assignment.) **(F)** GSEA comparing differentially expressed genes in isotype switched clusters to the ranked gene list of IIV-elicited H1-specific FcRL5^+^ over FcRL5^neg^ IgD^neg^ B cells. (ref 16) **(G)** UMAP projection shows areas of differential abundance (DA-seq, ref 23) as overrepresented in LAIV (blue, DA1) versus IIV inoculated (red, DA2) subjects. **(H-I)** Dot plot of genes that are predicted by DA-seq as over-expressed in DA1 **(H)** or DA2 **(I)**.

Next, to finely assign clusters to B cell subsets, we evaluated the clusters for frequency of somatic hypermutation [Fig 4B], isotype distribution [Fig 4C], and clonality [Fig 4D]. Since cluster 10 had few cells (<100), it was excluded from further analysis. We found cluster 7 cells to be relatively unmutated, largely of an IgM isotype and polyclonal and therefore assigned these cells as naïve B cells. In contrast, clusters 2 and 4 were found as almost completely isotype switched and significantly more somatically hypermutated than cluster 7 naïve B cells. Clusters 2 and 4 comprised expanded clones (defined as clones with >3 highly similar sequences). We found clusters 0, 1 and 8 to be isotype switched, polyclonal and relatively less mutated that clusters 2 and 4 cells. Cluster 5 comprised cells with a predominant IgM isotype. Although cluster 5 was less mutated than clusters 2 and 4, it comprised expanded clones, including a large clone with 107 highly similar sequences from a single IIV inoculated subjected. Cells in cluster 3 were predominantly isotype switched, comprised expanded clones but were significantly less mutated than cells in clusters 2 and 4.

We and others have shown that isotype-switched HA^+^ memory B cells after IIV can be classed into subsets with stem-like and effector memory functions.^16^ To determine if clusters that comprise largely isotype-switched cells (0,1, 2, 3, 4 and 8) were enriched in genes associated with stem-like or effector memory B cell functions, we performed GSEA against known stem-like and effector memory B cell genes that we previously described [Fig 4E].^16^ We found clusters 0, 1 and 8 as enriched with stem-like memory B cell genes. We found cluster 3 as enriched in effector memory genes. Clusters 2 and 4 did not significantly enrich with either effector or central memory genes. Instead, we found cluster 2 had down-regulated stem-like memory genes, *CCR7*, *BACH2*, and *CXCR5* but had not upregulated effector memory genes. Cluster 4 up-regulated a limited number of effector memory genes, like *BHLHE40*, *CXCR3*, and *TKT*. Collectively, our unsupervised analysis of HA^+^ IgD^neg^ B cells after LAIV or IIV indicate that these cells comprise several circulating B cell populations with potential stem-like or effector-like functions.

To start to understand differences between cells from LAIV inoculated versus IIV inoculated individuals, we colorized cells by assignment as IIV-elicited or LAIV-elicited and found areas in UMAP space wherein cells preferentially belonged to one vaccine platform over the other [Supplemental Fig 4D]. To test this possibility more rigorously, we applied a multiscale analytic approach, DA-seq^23^, that uses k-nearest neighbor clustering to define differentially abundant populations between two biological states. Our DA-seq analyses defined a region associated with cluster 3 as over-represented in patients who received LAIV (DA1) and a region associated with cluster 4 as over-represented in patients who received IIV (DA2) [Fig 4F]. Marker genes upregulated in DA1 include genes known to regulate NK cell cytotoxicity, *CST7*^24^ and *SLAMF7*,^25^ a gene that encodes proteoglycan protein with the macromolecular granzyme and perforin complex^26^, *SRGN*, genes that regulate effector cell functions and exhaustion programs, *ZEB2*^27^ and *TOX*^28^, as well as genes related to TNF-α signaling, *TNFRSF1B* and *TNIP2* [Fig 4G].^29^ Marker genes associated with DA2 include genes that regulate cell-cycle progression, *CDKN1A*^30^ and *SIK3*^31^, genes that regulate NFκB signaling, *NFKB1*^32^ and *NFKBID*^33^, and genes that regulate the unfolded protein response and endoplasmic reticulum stress, *ATF6*^34^ and *EIF2AK3*^35^ [Fig 4H]. Thus, and in keeping with our supervised bulk RNA-seq analyses, our unsupervised single cell data sets also suggest that LAIV and IIV elicit transcriptionally distinct circulating HA^+^ IgD^neg^ B cells.

### LAIV-elicited HA^+^ IgD^neg^ B cells Are Not Uniformly Recalled as Peripheral Plasmablasts after Antigen Re-challenge

To better understand functional distinctions between clusters 3 and 4 that are differentially represented in circulation after LAIV and IIV respectively, we evaluated their relative expression of previously defined canonical memory B cell TFs and transcriptional regulators (TRs) [Fig 5A]. While both clusters expressed near equivalent levels of the memory B cell TF, *HHEX*, suggesting a memory B cell fate commitment, we found a graded increase in the expression of the master effector cell TF *TBX21* with intermediate expression in cluster 4 and highest expression in cluster 3. Cluster 4 was distinguished from cluster 3 by expression of the TF *RUNX3* and *BHLHE40* that enforces effector memory programs and blocks terminal effector cell differentiation in CD8 T cells.^36, 37, 38, 39^ Cluster 3 was distinguished from cluster 4 by high expression of the terminal CD8 T cell effector cell TF, *ZEB2*.

**Figure 5.**
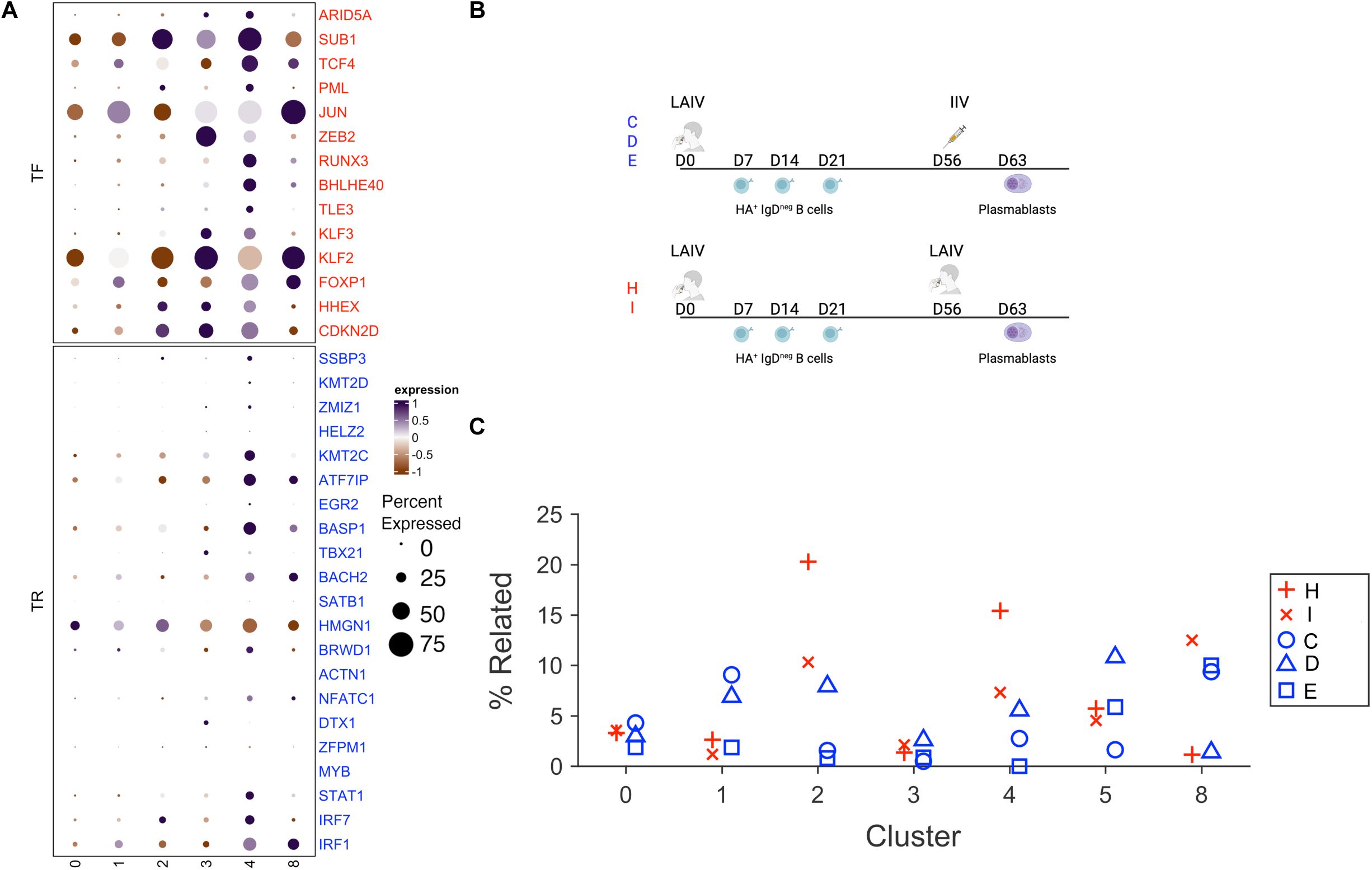
Recall potential of LAIV-elicited HA^+^IgD^neg^ B cells is heterogeneous. **(A)** Dot plot of transcription factors (TFs, colorized as red) and transcription regulators (TRs, colorized as blue) for isotype switched B cell clusters depicted in Figure 4A. **(B)** Schematic of immunization schedule for data represented in C. **(C)** Scatterplot shows percent clonal sharing between HA^+^ IgD^neg^ B cells by transcriptional cluster assignment (depicted in Figure 4A) and PBs after boost. Subjects boosted with IIV are colorized as red and subjects boosted with LAIV are colorized as blue.

Since cluster 3 over-expresses terminal cell differentiation marker, *ZEB2*, relative to cluster 4, we hypothesized that the recall potential of these clusters after antigen re-challenge may differ. To test this possibility, we boosted N = 3 LAIV-inoculated subjects with IIV 60 days later [Fig 5B], sort-purified IIV-elicited PBs and tested for clonal sharing between HA^+^IgD^neg^ B cells that circulate after LAIV prime and PBs that circulate after IIV boost. Shared clones were defined as 85% CDR3 nucleotide similarity and the same V, J and CDR3 length. Using these metrics, we found VDJ sequences from LAIV-elicited HA^+^ IgD^neg^ B cells assigned to cluster 3 were under-represented in the subsequent IIV-elicited PB repertoire, while VDJ sequences from LAIV-elicited HA^+^ IgD^neg^ B cells assigned to clusters 2 and 4 were over-represented in the subsequent IIV-elicited PB repertoire [Fig 5C].

We reasoned that this finding could be attributed to either (1) the possibility that cluster 3 cells function as terminal effectors and die after antigen re-challenge or (2) the possibility that certain LAIV-elicited cells have different systemic versus mucosal lymph node origins from IIV-elicited cells and may therefore be relatively less available for boost after IIV. In support of the second possibility, we found the rates of somatic hypermutation between LAIV and IIV-inoculated subjects to differ by transcriptional cluster assignment [Supplemental Fig 5A], suggesting that the lymph node precursors of cells that belong to the same transcriptional cluster may differ by vaccine platform.

Therefore, to test the second possibility more rigorously, we performed VDJ and GEX sequencing of the HA^+^ IgD^neg^ B cells that circulate at days 7, 14, and 21 after LAIV inoculation of one individual and tested for clonal sharing between transcriptionally discrete clusters over time [Supplemental Fig 5 B-C]. We identified the same transcriptionally discrete clusters that we observed previously [Fig 4A] at all weekly time points (*data not shown*). We found minimal clonal sharing between clusters 3 and other clusters at a single time point or over time after LAIV inoculation [Supplemental Fig 5B]. We also found that clusters 2 and 4 comprised expanded clones (>4 highly similar sequences) that were clonally related (defined as 85% CDR3 nucleotide similarity and identical V, J gene annotation) at day 14 and day 21 after immunization [Supplemental Fig 5C.] Collectively, these data support the possibility that the lymph node origins of cluster 3 may be unique from other LAIV-elicited HA^+^ IgD^neg^ B cells.

Therefore, we hypothesized that LAIV-elicited cluster 3 cells may be more efficiently recalled back into blood as PBs after mucosal boost with LAIV. To test this possibility, we boosted N =2 LAIV inoculated subjects with an additional dose of LAIV 56 days after prime [Fig 5B], sort-purified day 7 circulating PBs and evaluated for clonal sharing between HA^+^IgD^neg^ cells that circulate after LAIV prime and the PBs that circulate after LAIV boost. We could readily identify clones from clusters 2 and 4 after LAIV prime that were recalled as circulating PBs after LAIV boost [Fig 5C], suggesting that cells in these clusters are not terminally differentiated and can form PBs after either parenteral or mucosal antigen challenge. However, we could not identify clones from cluster 3 cells after LAIV prime that were recalled as circulating PBs even after mucosal LAIV boost. Together, these data suggest that in contrast to IIV, LAIV preferentially elicit a subset of circulating T-bet^+^ HA^+^ IgD^neg^ B cells that are marked by terminal effector cell transcriptional programs that cannot be efficiently boosted for immune protection as PBs after systemic or mucosal antigen re-challenge.

## Discussion

Traditionally defined by cell surface marker expression, human memory B cells are now realized to be heterogeneous by antigen-specificities and transcriptional programs.^40^ Memory B cells that express the transcription factor, T-bet, have been described after chronic infections, autoimmunity and immunization. A recent comparative analysis of T-bet^+^ B cells in various chronic infectious and autoimmune diseases demonstrate that the transcriptional properties and functional attributes of these cells can differ by disease state.^41^ Our study adds to this literature by presenting data in an immunization model that also demonstrate the transcriptional and functional heterogeneity of the circulating T-bet^+^ B cell compartment. We specifically show that mucosal inoculation with LAIV preferentially elicits circulating T-bet^+^ B cells that are characterized by expression of genes associated with terminal effector cell differentiation and function as terminally differentiated because they cannot be recalled as circulating PBs after antigen re-challenge. Finally, these cells appear to be clonotypically unique and less mutated than other LAIV-elicited HA^+^ IgD^neg^ B cells, suggesting that these cells may arise from different lymph node precursors.

Even if modest in magnitude, graded T-bet expression has been shown as determining differences in CD8 T cell fate with highest expression defining a short-lived effector cell fate and lower expression defining a memory precursor cell fate.^42^ Graded T-bet expression in CD8 T cells is established by cytokines, like IL-12, during T cell priming. Mechanistically, high T-bet expression promotes terminal CD8 T cell effector fate in concert with the TF *ZEB2*, which allows T-bet to bind to the promoter regions of terminal effector cell genes.^43^ Our B cell data echo these findings from CD8^+^ T cells and show that highest *TBX21* expression co-express *ZEB2* and associate with a terminal effector B cell fate after immunization. Future epigenetic studies should assess if *ZEB2* functions cooperatively with T-bet to promote gene expression of terminal effector genes in B cells as has been described in CD8 T cells. Cytokine signals that dictate these various B cell fate decisions after immunization should also be explored.

Notably, our findings derive from a cohort of LAIV-inoculated adult subjects with low baseline serum reactivity to vaccine antigen, who seroconverted after LAIV; this cohort is not necessarily representative of all LAIV-inoculated adults. Certain studies link seroconversion after LAIV with replication competent viral shedding shortly after live attenuated mucosal immunization.^44, 45, 46^ These data imply that our study cohort who seroconverted after LAIV may be distinguished by transient and productive mucosal infection after LAIV administration. Thus, our reported transcriptional differences between LAIV-elicited and IIV-elicited HA^+^ IgD^neg^ B cells may result from differences in the inoculum as replication competent (LAIV) versus replication incompetent (IIV) and may not necessarily result from differences in inoculum placement as mucosal over systemic.

Paired mucosal and systemic sampling after LAIV-inoculation of adults shows a decoupling of nasal and systemic Ab responses to vaccine with as many as 40% of adult subjects mounting a nasal Ab response alone.^47^ More recently, adenoid sampling of adults who did not sero-convert after LAIV has been reported and these adenoid samples demonstrate that even in the absence of systemic sero-conversion, LAIV elicits mucosal HA^+^ IgD^neg^ B cells as well as HA^+^ GC B cells.^48^ Thus, systemic and mucosal responses to LAIV may be separate and our finding that terminally differentiated HA^+^ IgD^neg^ B cells are enriched in the blood after LAIV may not indicate a similar enrichment of terminally differentiated HA^+^IgD^neg^ B cells in the mucosa.

Although a limitation of our study is that we do not directly link the systemic circulating HA^+^ IgD^neg^ B cell response to the quality of mucosal immune protection after LAIV, we do identify two circulating populations of HA^+^ IgD^neg^ B cells (clusters 2 and 4) that are clonally expanded with clones that can be repetitively sampled in the blood at days 14 and 21 after LAIV inoculation [Supplemental Fig 4B]. These data suggest clusters 2 and 4 may arise from proliferating mucosal HA^+^ GC B cell precursors that transiently re-circulate after LAIV inoculation. Intriguingly, lineages from cells in clusters 2 and 4 can be identified more often in the circulating PBs repertoire after mucosal over systemic antigen re-challenge, suggesting that cluster 2 and 4 cells may seed the mucosa as resident cells and thus become more available for boost after mucosal antigen encounter. In support of this possibility, cluster 4 upregulates TFs associated with successful mucosal tissue residence of CD8 T cells, *BHLHE40* and *RUNX3*.^49, 50^ Future studies should enroll larger cohorts of LAIV-inoculated healthy subjects in order to test if mucosal versus systemic inoculum boosts different circulating and mucosal cohorts of HA^+^ IgD^neg^ B cells as circulating or mucosal ASCs. Studies that directly assess clonal relationships, reactivities and breadth of neutralization between matched mucosal and circulating HA^+^ IgD^neg^ B cells are also required.

## Author Contributions

A.N. F.E.L. and T.D.R conceived the idea for the project and secured the initial funding. A.N. designed the experiments that were performed by A.N., E.Z., C.D.S., J.T.G, J.L. B.M. and F.Z. B cell tetramers were developed and produced by J.A. Human samples used in this study were obtained via the Alabama Vaccine Research Clinic, directed by P.A.G. Bioinformatic analyses were performed by A.F.R, C.D.S., E.I., and C.F. All other data was analyzed by A.N. A.N. wrote the manuscript and prepared final figures. Critical feedback on the project and manuscript was provided by R.S.H, R. D., P.M, V.E.N., A.F.R, T.D.R, F.E.L, P.A.G, C.D.S.

## Acknowledgements

We acknowledge Sergei Koralov for his thoughtful comments, edits and suggestions. We additionally acknowledge the members of the UAB Single Cell Core (Shanrun Liu) and Flow Cytometry Facility (Hanumanthu Sagar) for their technical assistance.

## Methods

### Human Subjects and Samples

The UAB Institutional Review Board approved all human study protocols. Subjects, who self-identified as healthy and having missed the prior season’s influenza vaccine, were recruited and consented through the Alabama Vaccine Research Clinic (AVRC). Subjects were enrolled to receive their first seasonal vaccine as the 2015-2016 Fluzone (Sanofi-Pasteur), the 2015-2016 Flumist (AstraZeneca), the 2018-2019 Flumist (AstraZeneca), the 2018-2019 Fluzone (Sanofi-Pasteur), the 2019-2020 Flumist (AstraZeneca) or the 2019-2020 Fluzone (Sanofi-Pasteur).

Blood was drawn on days 0, 7, 14, 21, and 120 days +/− 1 week from subjects enrolled in 2015-2016 and on days 7, 14, 21 from subjects who received all other vaccines. Subjects who received 2018-2019 Flumist were boosted (N = 3) with the 2018-2019 Fluzone 56 days later and blood was drawn at day 63 after boost. Subjects who received the 2019-2020 Flumist (N = 2) were boosted with an additional 2019-2020 Flumist 56 days later and blood was drawn at day 63 after boost. These prime boost studies were done as part of a UAB IRB approved clinical trial protocol NCT04080245.

### Lymphocyte and plasma isolation

Peripheral blood was drawn into K2-EDTA tubes (BD Bioscience). Peripheral blood mononuclear cells (PBMCs) and plasma were isolated by density gradient centrifugation over Lymphocyte Separation Medium (CellGro). Red blood cells were lysed with ammonium chloride solution (StemCell). Plasma and PBMCs were either used immediately for single cell sequencing or aliquoted and stored at −80°C for assays.

### Recombinant influenza HA protein production

Coding sequences (amino acids 18-524) of influenza HA ectodomains were synthesized (GeneArt, Regensburg, Germany) from vaccine antigen matched strains. HA ectodomains were cloned into the pCXpoly(+) vector that was modified with a 5’ human CD5 signal sequence and a 3’ GCN4 isoleucine zipper trimerization domain (GeneArt) that was followed by either a 6XHIS tag (HA-6XHIS construct) or an AviTag (HA-AviTag construct).^4^ The HA–6XHIS and HA–AviTag constructs for each HA were co–transfected using 293Fectin Transfection Reagent into FreeStyle™ 293–F Cells (ThermoFisher Scientific) at a 2:1 ratio, respectively. Transfected cells were cultured in FreeStyle 293 Expression Medium for 3 days and the supernatant was recovered by centrifugation. Recombinant HA molecules were purified by FPLC using a HisTrap HP Column (GE Healthcare) and eluted with 250 mM imidazole.

### Flow Cytometry

Single cell suspensions were blocked with 2% human serum and stained with Ab panels described in Supplemental Table 2A. 7AAD or LIVE/DEAD Fixable Dead Cell Stain Kits (Molecular Probes/ThermoFisher) were used to discriminate live cells. To eliminate non-specific HA tetramer binding, cells were treated at 37°C with 0.5U/ml neuraminidase (*C. perfringens*, Sigma) to remove sialic acid, and then were washed and stained with HA tetramers. Intracellular proteins were detected by staining with Abs specific for cell surface markers, fixing the cells in 10% neutral buffered formalin solution (Sigma), and then staining the permeabilized cells (0.1% IGEPAL (Sigma) with intracellular antibodies. Stained cells were acquired for analysis with either a FACSCanto II (BD Bioscience) or the BD Symphony flow cytometer (Invitrogen, ThermoFisher). FlowJo v9.9.3 or FlowJo v10.2 were used for analysis.

### Cell Sorting

B cell subsets were sort-purified for bulk RNA-seq or single cell VDJ and GEX analyses with a FACSAria (BD Biosciences) in the UAB Comprehensive Flow Cytometry Core. Supplemental Table 2B shows flow panels used for subset sort-purifications.

### RNA-seq library preparation and data analyses

B cell subsets were sorted directly into RLT buffer (Qiagen) with 1% mercaptoethanol and then snap-frozen in LN_2_. RNA was extracted using the QuickRNA Micro Prep Kit (Zymo) and cDNA was prepared using SMART-seq v4 cDNA synthesis kit (Takara). Sequencing libraries were constructed using 200 pg cDNA as input for the NexteraXT kit with NexteraXT indexing primers (Illumina). Libraries were quality assessed, pooled and sequenced using 75 bp paired-end chemistry on a NextSeq500 at the UAB Helfin Genomics Core. Sequencing reads were mapped to the hg38 version of the human genome using STAR ^51^ with the default settings and the UCSC KnownGene table as a reference transcriptome. Reads overlapping exons were tabulated using the GenomicRanges ^52^ package in R/Bioconductor. Genes expressed at 3 reads per million or more in all samples from one group were considered detected and used as input for edgeR v3.24.3 ^53^ to identify DEG. P-values were FDR corrected using the Benjamin-Hochberg method ^54^ with genes of an FDR <0.05 and an absolute log_2_ FC >1 considered significant. Expression data was normalized to reads per kilobase per million mapped reads (RPKM). Samples included either 3 LAIV samples (D14 timepoint) or 6 IIV (D14 timepoint) biological replicates per B cell subset.

### Single Cell RNA-library preparation and sequencing

Five LAIV inoculated subjects and five IIV inoculated subjects were enrolled into study and had blood drawn at days 7, 14, 21 after immunization. B cells were negative selected using column purification (EasySep Pan-B cell purification kit, StemCell Technologies), treated with neuraminidase, stained with HA tetramers and Abs conjugated with TotalSeq C oligomer hashtags [Supplemental Table 3]. Single cell suspensions were applied to the 10XGenomics workflow for cell capture, scRNA gene expression (GEX), BCR and Feature Barcoding library preparation using the Chromium Single Cell 5’ Library and Gel Bead Kit (Nest GEM 5’ Kit v2) as well as the Single Cell 5’ Feature Barcode Library Kit (10XGenomics), following manufacturer’s instructions.

### Single-cell transcriptome and repertoire profiling

Raw sequence reads were processed using CellRanger (version 7.1) with multi-pipeline and default parameter settings used for demultiplexing and quantifying gene expression and assembly of BCR sequences. Reference datasets used for GEX and VDJ annotation included refdata-gex-GRCh38-2024-A and refdata-cellranger-vdj-GRCh38-alts-ensembl-7.1.0 respectively (10x Genomics). Cellranger output was analyzed in R (version 4.4.1) using Seurat package (version 5.1.0). Cellranger count data of the enrolled subjects were log normalized, transformed and variable genes were detected with uniform manifold approximation and projection (UMAP) performed in default parameter settings, using Seurat with 15 principal components. Ambient RNA levels were detected and accounted for using SoupX.^55^ Doublet formation was estimated and cells identified as likely being doublets were removed using DoubletFinder.^56^ Transcriptionally similar clusters were identified with shared nearest neighbor (SNN) modularity optimization with a resolution of 0.8. High confidence V_H_ consensus sequences were analyzed using IMGT High-Vquest (version 3.6.3; reference release 202030-2) to retrieve VDJ annotation and nucleotide CDR3 sequence. Corresponding VDJ sequence annotation data was mapped back to known GEX barcode data for further analysis in Seurat.

### GSEA on single cell dat

(cluster 0, 1, 2, 3, 4, 8 vs FcRL5^+^ over FcRL5^neg^ differentially expressed genes) was performed using the parameters 0.1 min pct, padj <0.05.

### HA ELISAs

Plasma from vaccinated samples were serially diluted on EIA/RIA ELISA plates (Costar) that were previously coated with recombinant HA. HA-specific IgG Abs from the samples were detected using HRP-conjugated anti-human IgG secondary Abs (Jackson ImmunoResearch) and were developed using ABTS with acid stop. Absorbance was measured at 415nm using a SpectraMaxM2 (Molecular Devices). All samples were tested against the same reference standard and 50% endpoint titers were determined.

### Statistical Analyses

Comparisons between two matched groups (e.g. PBs at day 0 and day 7 from each subject) were performed with the Wilcoxson matched pairs signed rank test for non-normally distributed variables. Comparisons between unmatched groups was performed using the Mann-Whitney test. Strength and direction of association between two variables measures was performed using the D’Agostino-Pearson normality test followed by or Spearman correlation test. Data were considered significant when *P* ≤ 0.05. GraphPad Prism version 7.0a or 8.0 software (GraphPad) was used for analysis.

### Other bioinformatics analyses and visualizations

Scatterplots plots and other visualizations were created using Matlab (R2024b, The Mathworks Inc., Natick MA) or R v4.41.

---

**Supplementary Table 1.** Demographics of 2015-2016 Influenza Vaccinated Cohort.

|  |  |  |
| --- | --- | --- |
| **Characteristic** | **Vaccine Modality** |  |
|  | IIV (n = 19) | LAIV (n = 17) |
| Age years, mean (SD) | 30 (6) | 31(10) |
| Gender |  |  |
| % Male | 57.8% | 23.8% |
| % Female | 42.2% | 76.5% |
| Race |  |  |
| % Caucasian | 63.1% | 41.1% |
| % African-American | 36.8% | 41.1% |
| % Asian | 0% | 11.7% |
| % Other | 0% | 5.8% |

**Supplemental Table 2A.**
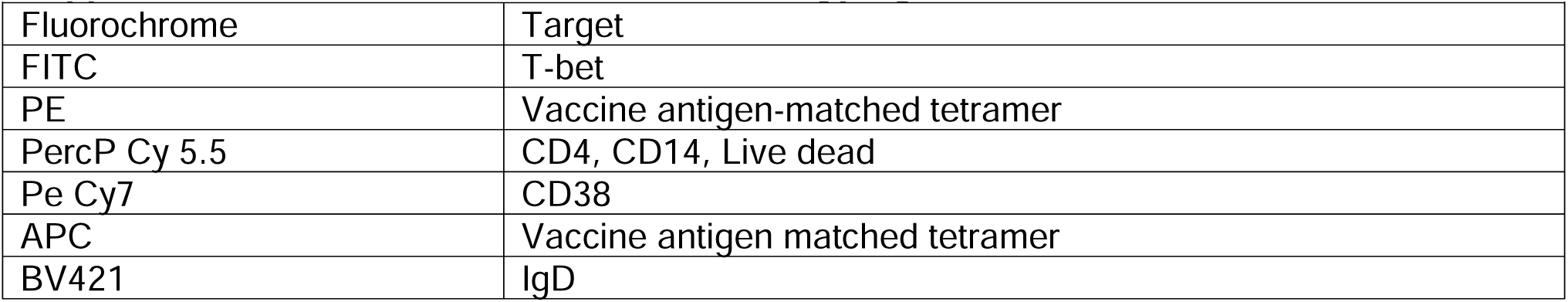
Flow Panels for Phenotyping.

| **Supplemental Table 2A. Flow Panels for Phenotyping** |  |
| --- | --- |
| Fluorochrome | Target |
| FITC | T-bet |
| PE | Vaccine antigen-matched tetramer |
| PercP Cy 5.5 | CD4, CD14, Live dead |
| Pe Cy7 | CD38 |
| APC | Vaccine antigen matched tetramer |
| BV421 | IgD |

**Supplemental Table 2B.** Flow Sort Panels for Bulk RNA-seq and Single Cell Sequencing.

|  |  |
| --- | --- |
| Fluorochrome | Target |
| FITC | IgD |
| PE | Vaccine antigen-matched tetramer |
| PercP Cy 5.5 | CD4, CD14, Live dead |
| Pe Cy7 | CD38 |
| APC | Vaccine antigen matched tetramer |

**Supplemental Table 3** Related to Figure 3. Bulk RNA-seq data from sort-purified HA^+^ IgD^neg^ B cells at day 14 after immunization.

**Tab 1**. RNA-seq analysis of sort-purified B cell subsets from immunized subjects reported as RPKM values for each gene. Log_2_FC, p values and FDR values are provided for each pairwise comparison.

**Tab 2**. Description of gene sets used for GSEA analyses including source of gene sets and list of genes.

**Tab 3**. DEG from LAIV-elicited FcRL5^+^ and FcRL5^neg^ IgD^neg^ B cell RNA-seq data sets (derived from Tab 1) and published data sets in ref 16 (Tab 2 for list) and Hallmark data sets. Significant genes are indicated as “1” and non-significant as “0” in the comparator gene set.

**Supplemental Table 4**. Related to Figures 4 and 5 Single cell sequencing data. Transcriptional and repertoire data from single cell sorted HA^+^ IgD^neg^ B cells from N = 5 LAIV vaccinated healthy subjects and N = 5 IIV vaccinated healthy subjects.

Tab 1 unique DEGs within clusters,

Tab 2 GSEA statistics,

Tab 3 oligomer hashtags used

Tab4 mutation frequencies

**Supplemental Figure 1** (accompanying data in Figure 1). **(A)** Gating strategy for B cells, IgD^neg^ B cells and PBs **(B)** Gating strategy for cTfh cells **(C-F)** 19 subjects received 2015-2016 LAIV and 17 subjects received the 2015-2016 IIV. **(C)** IgG antibody titer (aribtrary units, AU) against the Ca H1N1 strain in LAIV inoculated subjects at day 0 and day 120 after immunization **(D)** Percent of Ca09 H1-binding IgD^neg^ B cells that circulate after IIV (circle) and LAIV (square) inoculation.

**Supplemental Figure 2** (accompanying data in Figure 2). FACS plots from a single subject immunized with LAIV in 2015-2016. **(A)** T-bet expression in live IgD^neg^ B cells H3^+^ cells in T-bet^neg^ **(B)** and T-bet^+^ **(C)** gates **(D-E)** Correlation between the fold change in H3-IgG Ab at days 0 and 120 and the magnitude of **(D)** PB cells or **(E)** cTfh cells.

**Supplemental Figure 3** (accompanying data in Figure 3). **(A)** Bar plot of Normalized Enrichment Score for each Hallmark gene set is shown. Significant bar plots (q <0.05) are colorized purple.

**Supplemental Figure 4**. (accompanying data in Figure 4). Data refers to clusters depicted in Figure 3A. **(A)** Bar plot depicting cell numbers per cluster. **(B)** Bar plot depicting percentage of cells by participant assignment (1-10) in clusters. **(C)** Box plot showing average ROGUE score (ref 22) by cluster assignment. **(D)** UMAP colorized by cell assignment as arising from IIV-inoculated subject (pink) or LAIV-inoculated subject (blue).

**Supplemental Figure 5**. (accompanying data in Figure 5) **(A)** Violin plot shows mutation rates by cluster assignment (see Figure 4A for UMAP) for LAIV versus IIV inoculated subjects ****p <0.0001Wilcoxon rank sum test**(B)** Heat map depicts percent clonal sharing across clusters at an individual time point and at weekly time points after LAIV inoculation of N =1 healthy subject. Gray diagonal boxes list total numbers of clones in each cluster by time point. **(C)** Bubble plot of shared clones (sequences > 4) grouped by day after immunization and cluster assignment from an LAIV inoculated subject.

