## Supplemental Figures for "Terminally Differentiated Influenza-Specific Effector Memory B Cells Circulate after Live Attenuated Influenza Vaccination"

#### Supplemental Figure 1

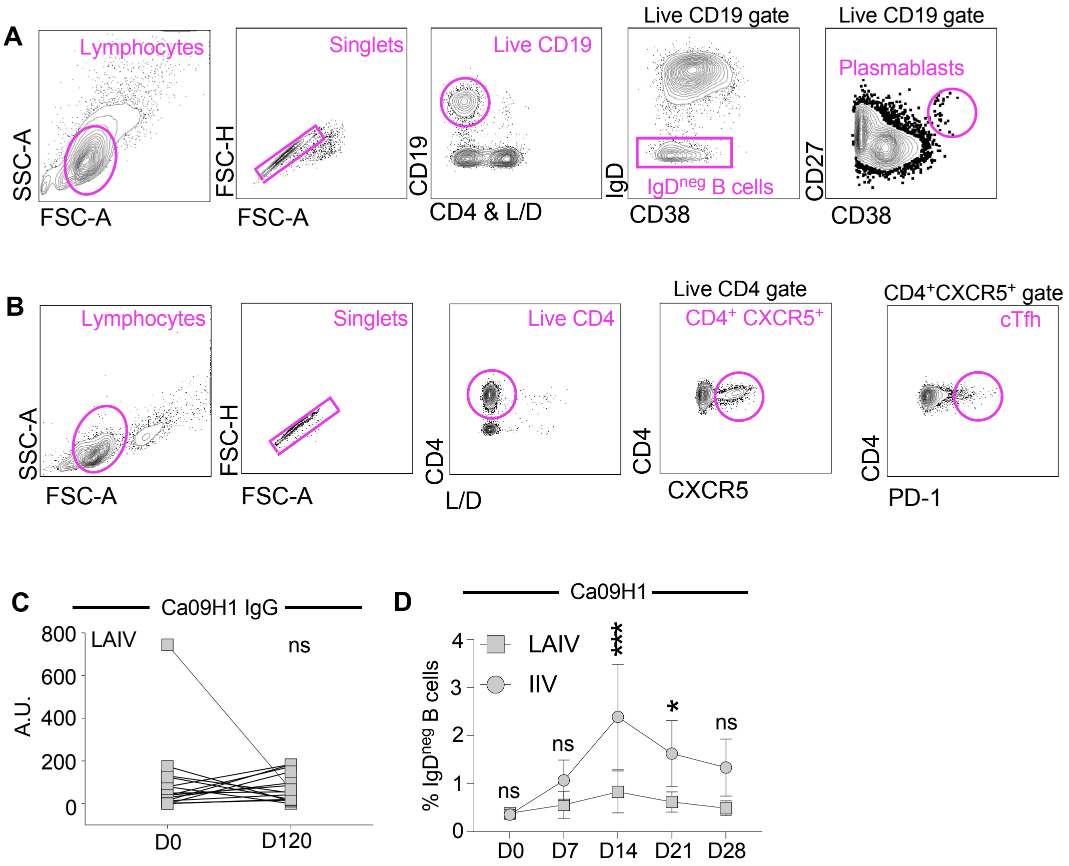

**Supplemental Figure 2**

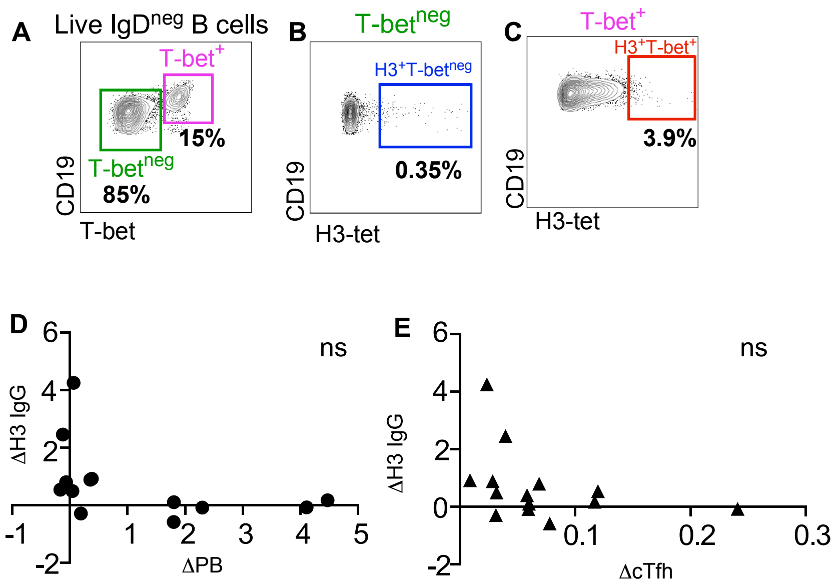

### Supplemental Figure 3

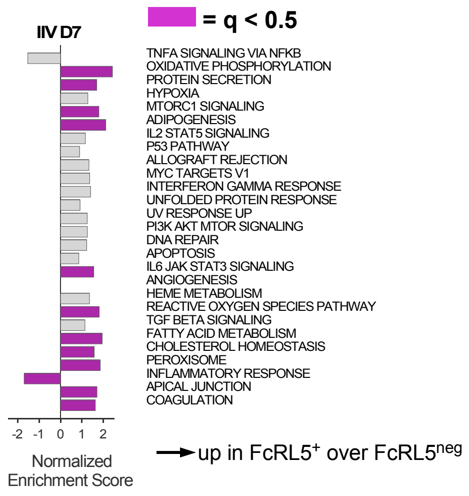

**Supplemental Figure 4**

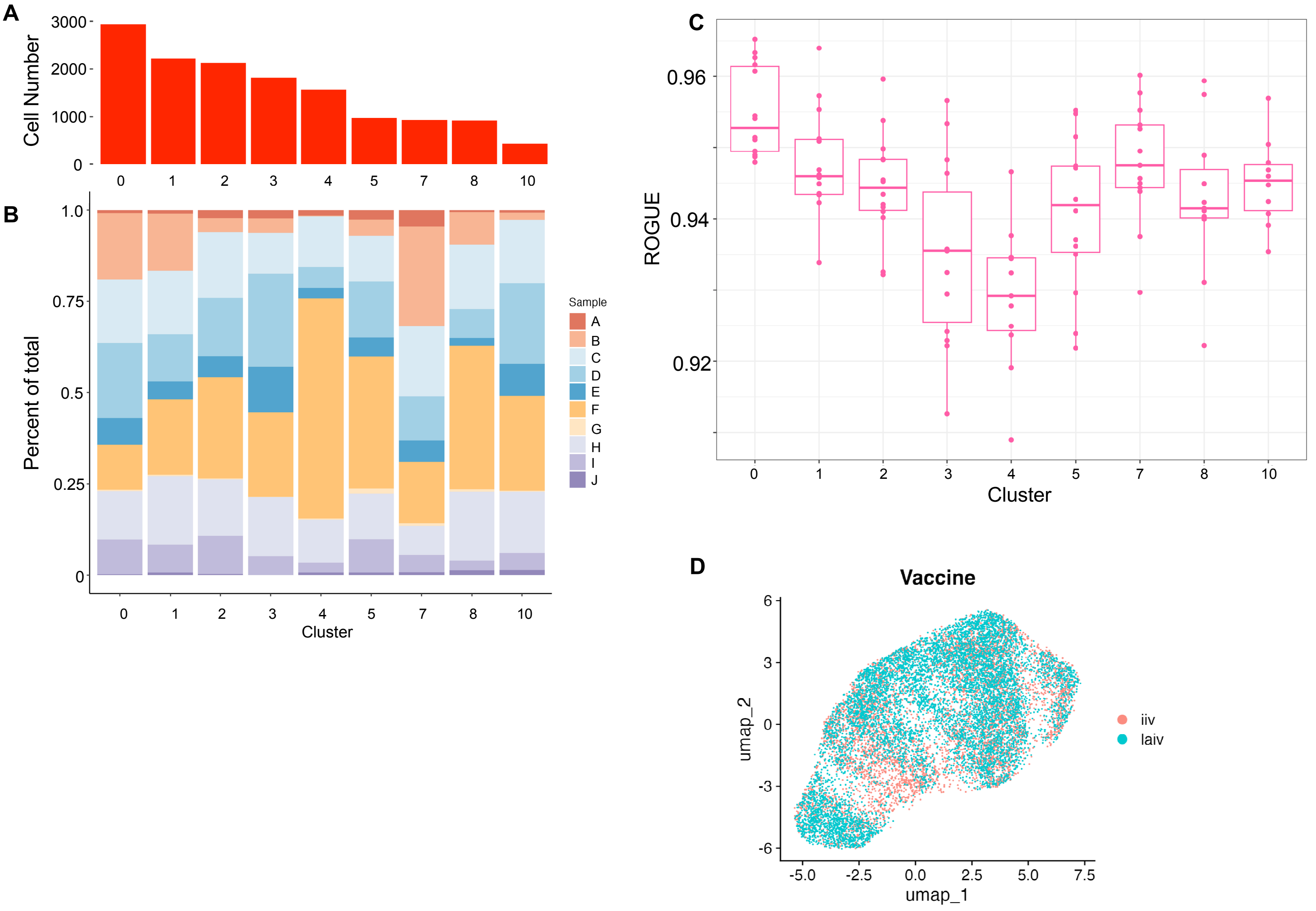

### Supplemental Figure 5

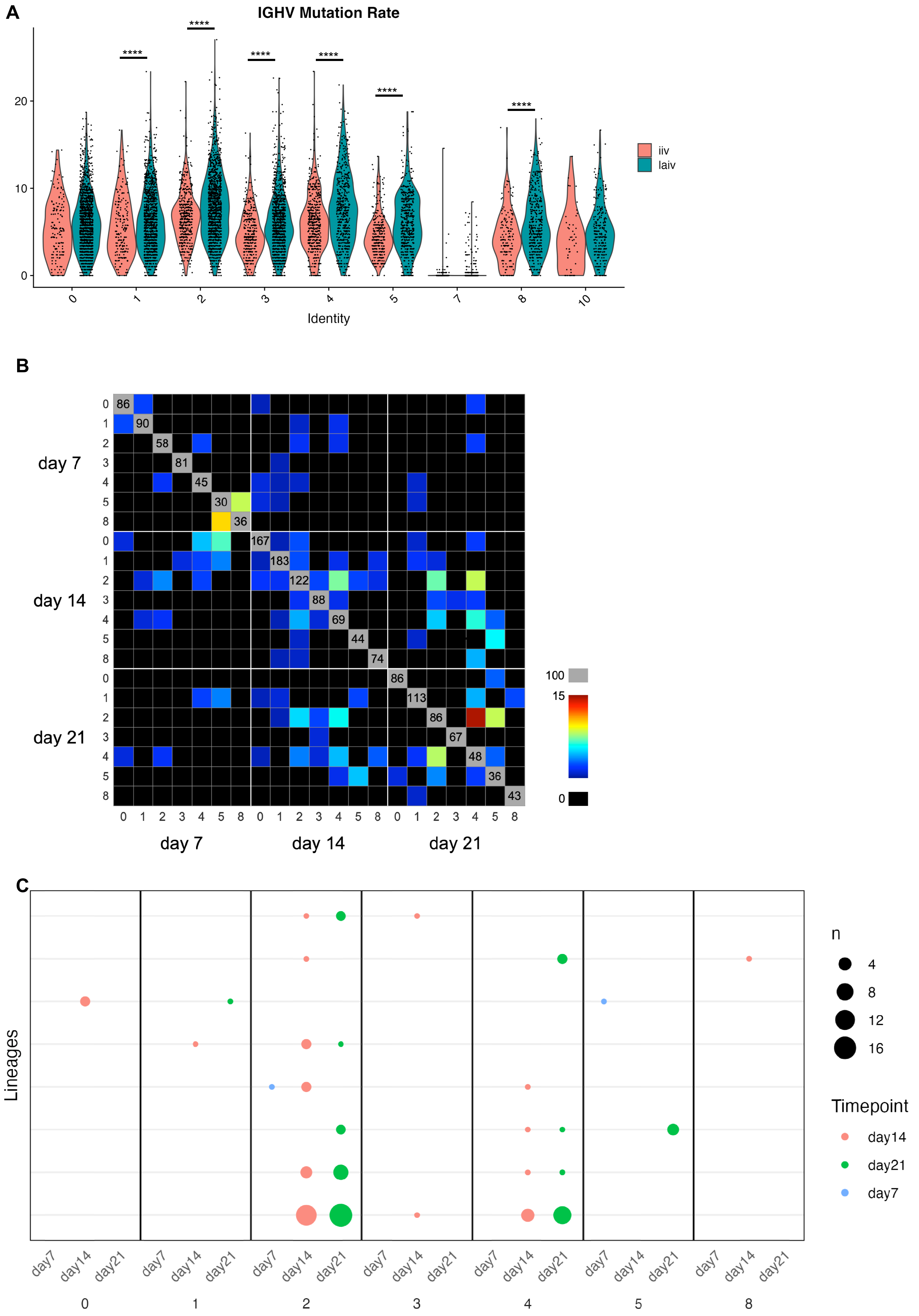
